# Host immune responses after suprachoroidal delivery of AAV8 in nonhuman primate eyes

**DOI:** 10.1101/2020.09.25.313676

**Authors:** Sook Hyun Chung, Iris Natalie Mollhoff, Alaknanda Mishra, Tzu-Ni Sin, Taylor Ngo, Thomas Ciulla, Paul Sieving, Sara M. Thomasy, Glenn Yiu

**Author notes:** Corresponding author Glenn Yiu, M.D., PhD., Department of Ophthalmology & Vision Science, University of California Davis, Address: 4860 Y St., Suite 2400 Sacramento, CA, 95817.

## Abstract

The suprachoroid is a potential space located between the sclera and choroid of the eye which provides a novel route for ocular drug or viral vector delivery. Suprachoroidal injection of AAV8 using transscleral microneedles enables widespread transgene expression in eyes of nonhuman primates, but may cause intraocular inflammation. We characterized the host humoral and cellular immune responses after suprachoroidal delivery of AAV8 expressing green fluorescent protein (GFP) in rhesus macaques, and found that it can induce a mild chorioretinitis that resolves after systemic corticosteroid administration, with recovery of photoreceptor morphology but persistent immune cell infiltration after 3 months. Suprachoroidal AAV8 triggered B-cell and T-cell responses against GFP, but only mild antibody responses to the viral capsid as compared to intravitreal injections of the same vector and dose. Systemic biodistribution studies showed lower AAV8 levels in liver and spleen after suprachoroidal injection compared with intravitreal delivery. Our findings suggest that suprachoroidal AAV8 primarily triggers host immune responses to GFP, likely due to sustained transgene expression in scleral fibroblasts outside the blood-retinal barrier, but elicits less humoral immune reactivity to the viral capsid than intravitreal delivery due to lower egress into systemic circulation. Thus, suprachoroidal AAV delivery of human transgenes may have significant translational potential for retinal gene therapy.

## Introduction

The first approved ocular gene therapy for treating biallelic RPE65 mutation-associated retinal dystrophy, Leber’s Congenital Amaurosis, has generated much enthusiasm for the use of adeno-associated viruses (AAVs) as vectors for retinal gene delivery.^1–6^ Recombinant AAVs are highly effective vectors for gene delivery due to their ability to transduce a wide variety of retinal cell types and relative safety given their nonpathogenic and non-integrating nature.^7^ However, although AAV vectors are much less immunogenic than adenoviruses, host immune responses triggered by the viral vector or transgene product can limit the effectiveness of the treatment.^8,9^ Humoral immune responses from neutralizing antibodies (NAbs) produced by B-cells can inhibit vector transduction. These antibodies may arise from prior exposure to wild-type AAV causing pre-existing immunity, or be triggered by therapeutic vector administration which prevents or suppresses further transduction.^10–13^ Also, cell-mediated immune responses from cytokine-secreting T-cells can directly destroy transduced cells.^14^ Together, host humoral and cellular immune responses contribute to eliminating vectors and transduced cells, thus limiting the therapeutic effect.

Although the eye has been considered to be an immunologically-protected space,^15^ the immunogenicity of AAV-mediated gene transfer in the eye varies with the route of administration. Subretinal injections, which entail a needle puncture through the neurosensory retina, enables efficient transduction of multiple cell types including photoreceptors and the underlying retinal pigment epithelium (RPE), and triggers minimal humoral immune responses.^16,17^ However, the procedure requires complex vitrectomy surgery and the therapeutic effect is limited to the area of the injected fluid bleb. Intravitreal injections can be easily performed in an outpatient clinical setting, and newer generations of AAV can overcome the internal limiting membrane (ILM) barrier to transduce deeper retinal layers.^18,19^ But unlike subretinal injections, intravitreal delivery triggers more pronounced humoral and cellular responses against the AAV capsid, occasionally to levels matching systemic administration.^13,20^

We and others have recently described a novel mode of ocular gene delivery by injecting AAV into the suprachoroidal space, which is located between the scleral wall and the choroidal vasculature of the eye.^21,22^ Although this potential space is barely detectable under physiologic conditions,^23,24^ suprachoroidal injection of compounds using transscleral microneedles expands the suprachoroidal space as seen on *in vivo* imaging,^25,26^ enabling targeted drug delivery to retinal and choroidal tissues while minimizing adverse effects on anterior segment structures.^27–31^ Suprachoroidal injection of a triamcinolone acetonide suspension using these microneedles has been effective in treating macular edema from noninfectious uveitis in human clinical trials.^32^

Using nonhuman primates (NHPs), we found that suprachoroidal injection of AAV8 using transscleral microneedles enables widespread, peripheral transduction of mostly RPE cells. By contrast, subretinal injection of AAV8 transduced outer retinal cells including photoreceptors and RPEs, but was limited to the injection site.^21^ Since the suprachoroidal space is located outside the blood-retinal barrier, we also investigated the inflammatory response in retinal and choroidal tissues, and found a greater degree of local immune cell infiltration after suprachoroidal delivery of AAV8 compared with subretinal or intravitreal injections. Interestingly, we found that intravitreal AAV8 triggered more serum NAbs than the other modes of injection, likely due to differences in the pharmacokinetics and biodistribution of the different modes of ocular AAV delivery.

In this ancillary study, we explore in detail the host humoral and cellular immune responses to suprachoroidal AAV8 in these rhesus macaques. Like humans, NHPs are natural hosts for wild-type AAV and develop immune conversions to subclinical infection, making them an excellent animal model for predicting host immune responses to AAV vectors in humans. We found that suprachoroidal injection of AAV8 expressing green fluorescent protein (GFP) can elicit a transient chorioretinitis that clinically resolves after systemic corticosteroid administration, with recovery of photoreceptor morphology despite some persistence of immune cell infiltration over 3 months. Suprachoroidal injections trigger both B-cell and T-cell responses against the GFP transgene product, whereas the response against AAV8 capsid was minimal compared with intravitreal injections. Systemic biodistribution assays showed limited presence of the AAV8 in the liver and spleen after suprachoroidal injections as compared with intravitreal delivery. As suprachoroidal injection of AAV is currently under evaluation for retinal gene therapy in human clinical trials, our results provide an important, clinically-relevant, and unique exploration of host immune responses from viral gene delivery to different ocular compartments surrounding the blood-retinal barrier.

## Results

### Study design and clinical course

Experiments to evaluate the transduction efficacy, pattern, durability, and cell-type specificity of suprachoroidal AAV8 injections in rhesus macaques using transscleral microneedles have been previously described.^21^ Briefly, we identified 5 animals between age 4-10 years with no pre-existing NAbs against AAV8, and injected both eyes with NHP-grade AAV8 that expresses enhanced GFP under a cytomegalovirus (CMV) promoter at 7 × 10^11^ vg/eye (low dose) or 7 × 10^12^ vg/eye (high dose), using either a 700-μm long 30-gauge microneedle (Clearside Biomedical, Alpharetta, GA, USA) for suprachoroidal or transscleral subretinal injection, or a 0.5-inch-long 30-gauge conventional needle for intravitreal injection (Supplementary Table 1). Of these, two animals received suprachoroidal AAV8 in both eyes (Rhesus 01 with 7 × 10^11^ vg/eye and Rhesus 02 with 7 × 10^12^ vg/eye), two animals (Rhesus 03 and 04) received suprachoroidal injection of AAV8 in one eye (7 × 10^12^ vg/eye) and subretinal delivery of AAV8 in the contralateral eye (7 × 10^12^ vg/eye), and the last animal (Rhesus 05) received intravitreal injection of AAV8 in both eyes (7 × 10^12^ vg/eye). After 1 month, suprachoroidal delivery of high-dose AAV8 produced diffuse, peripheral, and circumferential GFP fluorescence with a punctate pattern of expression (Figure 1A). By comparison, subretinal AAV8 resulted in a focal area of intense GFP expression (Figure 1B), while intravitreal AAV8 only produced a small peripapillary area of faint expression at the same high dose (Figure 1C). Suprachoroidal delivery of low-dose AAV8 (7 × 10^11^ vg/eye) did not produce any detectable transgene expression on fundus fluorescence imaging.

**[Figure 1].**
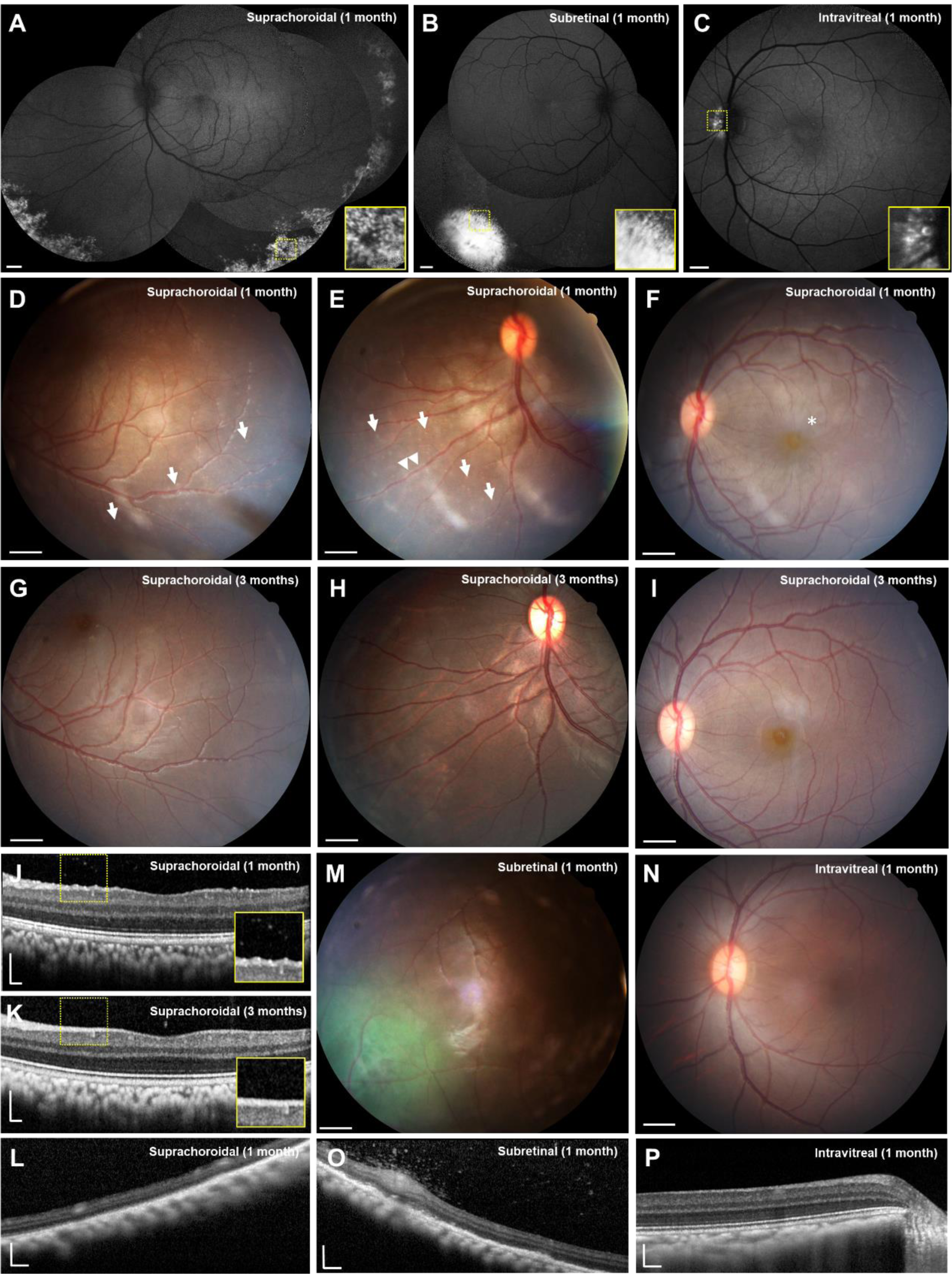
Multimodal ocular imaging after suprachoroidal, subretinal, or intravitreal injections of AAV8 to express enhanced GFP in NHP eyes. **(A-C)** Representative scanning laser ophthalmoscopy (SLO) montages and magnified insets of the yellow-dashed regions show different patterns of GFP transgene expression at 1 month after suprachoroidal (A), subretinal (B), and intravitreal (C) AAV delivery. **(D-I)** Representative color fundus photographs demonstrate punctate spots (arrows), perivascular sheathing (arrowheads), and radial macular striae (asterisk) that are observed after suprachoroidal AAV8 injections at 1 month (D-F), but resolved by 3 months (G-I), consistent with a transient chorioretinitis and vasculitis. **(J-L)** Representative spectral-domain optical coherence tomography (SD-OCT) images and magnified insets of the yellow-dashed regions reveal hyperreflective foci seen after suprachoroidal AAV8 at 1 month (J) but not at 3 months (K) or in peripheral retina (L). **(M-N)** Fundus photographs of macaque eyes demonstrate GFP fluorescence after subretinal AAV8 (M), and no clear inflammation after subretinal or intravitreal AAV8 (N). **(O-P)** SD-OCT images showed that subretinal AAV8 also induced cellular extravasation from retinal vessels suggestive of a localized vasculitis (O), but not after intravitreal injections (P). Scale bars, 1 mm for SLO images and fundus photos; 200 µm for SD-OCT images.

Although most of the animals did not exhibit significant anterior chamber (AC) or vitreous inflammation throughout the study, Rhesus 02 developed mild 2+ AC cell based on Standardization of Uveitis Nomenclature (SUN) criteria at 2 weeks requiring treatment with oral prednisone (1 mg/kg) for 2 weeks with subsequent resolution of the AC cell by month 1. At 1 month, this animal also demonstrated a peripheral chorioretinitis with small, punctate spots (Figure 1D), some perivascular sheathing (Figure 1E), and radial retinal striae in the macular region without significant macular edema (Figure 1F), which all appear resolved by month 3 (Figures 1G-1I). Spectral domain-optical coherence tomography (SD-OCT) imaging showed fine, hyperreflective foci in the vitreous and retinal surface at 1 month (Figure 1J) indicating subclinical vitritis not readily seen on funduscopic examination, which also resolved after 3 months (Figure 1K). We did not note significant vitreous cell in the peripheral regions of the transduced retina in Rhesus 02, or in any other animals after suprachoroidal delivery of AAV (Figure 1L). Eyes that received subretinal AAV8 showed localized vascular dilation and perivascular hyperreflective foci in the vitreous in the most intense regions of GFP expression (Figure 1M and 1O), indicating localized vasculitis and subclinical vitritis in these animals. Eyes that received intravitreal injection of AAV8 showed no detectable vitritis, chorioretinitis, or vasculitis, even in the small peripapillary region of transduction (Figures 1N and 1P). Thus, suprachoroidal injection of AAV8 may trigger an anterior uveitis, peripheral chorioretinitis, and mild vitritis that all resolve with oral corticosteroid treatment over 2 weeks. Subretinal AAV8 can also trigger mild, localized vasculitis and vitritis in the area of transduction, while intravitreal AAV8 exhibit poor transduction but showed no detectable intraocular inflammation.

### Local inflammatory responses after suprachoroidal AAV8

We previously found that suprachoroidal AAV8 injection elicited greater local infiltration of inflammatory cells than transscleral subretinal or intravitreal injections at 1 month post-injection.^21^ In this study, we further characterize the local inflammation using immunohistochemistry at 2 and 3 months after suprachoroidal AAV8 delivery (Figure 2). GFP transgene expression was detectable in both RPE and scleral tissues at 1 month, but only persisted in the sclera at months 2 and 3, appearing mostly in spindle-shaped cells that resemble scleral fibroblasts. The GFP expression in the sclera was not visible on live fundus imaging likely due to blockage of the fluorescence by the darkly-pigmented RPE and uvea in rhesus macaques.^33^ Local infiltration of ionized calcium-binding adaptor-1 (Iba1)+ microglia and macrophages (Figures 2A-2E), CD45+ leukocytes (Figures 2F-2J), CD8+ cytotoxic T cells (Figures 2K-2O), as well as reactive gliosis as shown by glial fibrillary acidic protein (GFAP) staining (Figures 2P-2T), were detected through month 3 as compared to uninjected control animals. Interestingly, the outer retinal layers and RPE architecture appeared partly restored at month 3 in the animal that received systemic corticosteroids (Figures 2U-2Y). The animal that received low-dose suprachoroidal AAV8 injections also demonstrated GFP expression in the sclera, and exhibited a similar degree of local inflammatory responses at month 3 (Figures 2D, 2I, 2N, 2S, 2X).

**[Figure 2].**
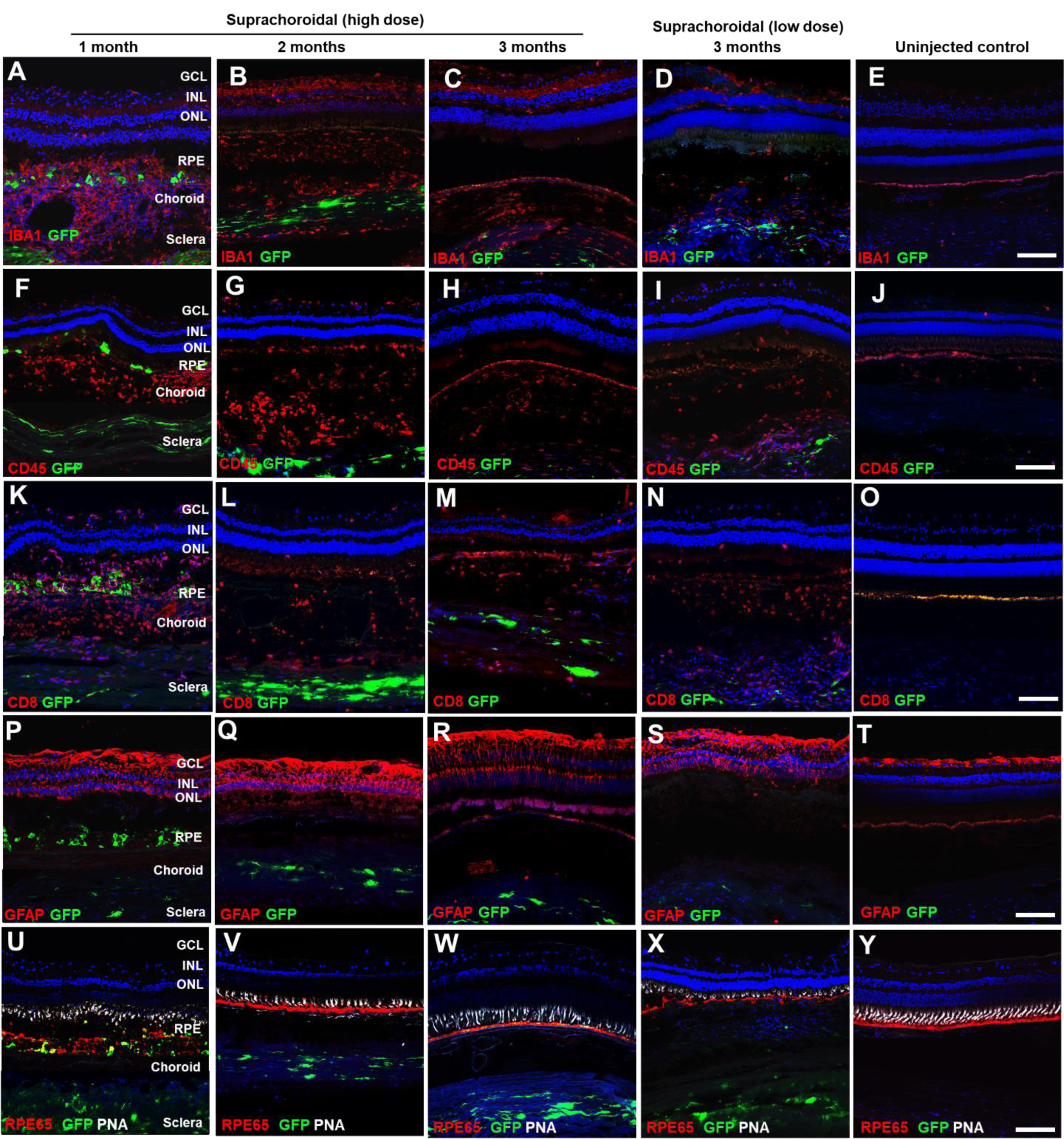
Local immune cell infiltration after suprachoroidal delivery of AAV8 in NHP eyes. **(A-Y)** Confocal fluorescence images of GFP transgene expression (green) co-immunostained with antibodies to IBA-1+ microglial cells (A-E), CD45+ leukocytes (F-J), CD8+ cytotoxic T-cells (K-O), and GFAP+ reactive gliosis (P-T), as well as RPE65 (red) to label RPE cells and peanut agglutinin (PNA, white) to label cone photoreceptor inner/out segments, along with DAPI (blue) to label cell nuclei in eyes at 1 month (A,F,K,P,U), 2 months (B,G,L,Q,V), and 3 months (C,H,M,R,W) after high-dose or low-dose (D,I,N,S,X) suprachoroidal AAV8 injections, as compared to uninjected control eyes (E,J,O,T,V). Abbreviations: GCL, ganglion cell layers; INL, inner nuclear layer; ONL, outer nuclear layer; RPE, retinal pigment epithelium. Scale bars: 100µm.

### Humoral immune responses after suprachoroidal AAV8

To evaluate humoral immune response from B-cells, we employed a sandwich enzyme-linked immunosorbent assay (ELISA) to measure serum binding antibodies against the AAV8 capsid or GFP transgene product after suprachoroidal or intravitreal delivery of the AAV8 vector (Figures 3A and 3B). Most of the animals that received suprachoroidal AAV8 developed minimal antibody responses against the viral capsid, whereas the animal that received intravitreal AAV8 exhibited higher anti-AAV8 antibody levels within 1 month (Figure 3A). These results are consistent with our prior study which demonstrated higher concentrations of serum NAbs from intravitreal than suprachoroidal or subretinal AAV8 as measured using an *in vitro* transduction inhibition assay.^21^ By contrast, only animals that received suprachoroidal AAV8 developed anti-GFP antibodies, which reached the highest levels at month 3, while the animal that received only intravitreal AAV8 did not (Figure 3B). As Rhesus 02 received high-dose suprachoroidal AAV8 in both eyes, we further validated the humoral response to GFP by performing flow cytometry on peripheral blood mononuclear cells (PBMCs) collected from the serum of this animal, and found expansion of GFP-responsive plasma B-cells (CD19-,CD27+,CD38+, HLADRlow) after suprachoroidal AAV8 injection (Figure 3C, Supplementary Figure 1) which likely accounts for the greater production of systemic anti-GFP antibodies. Interestingly, the animal that received low-dose AAV8 (Rhesus 01) developed similar concentrations of anti-GFP antibodies (Figure 3B). Together, these findings suggest that although intravitreal AAV8 produces an earlier and more robust humoral response to the viral capsid, suprachoroidal delivery triggers greater antibody responses to GFP, possibly due to exposure of GFP-expressing scleral fibroblasts to systemic immune surveillance, given their location outside the blood-retinal barrier.

**[Figure 3].**
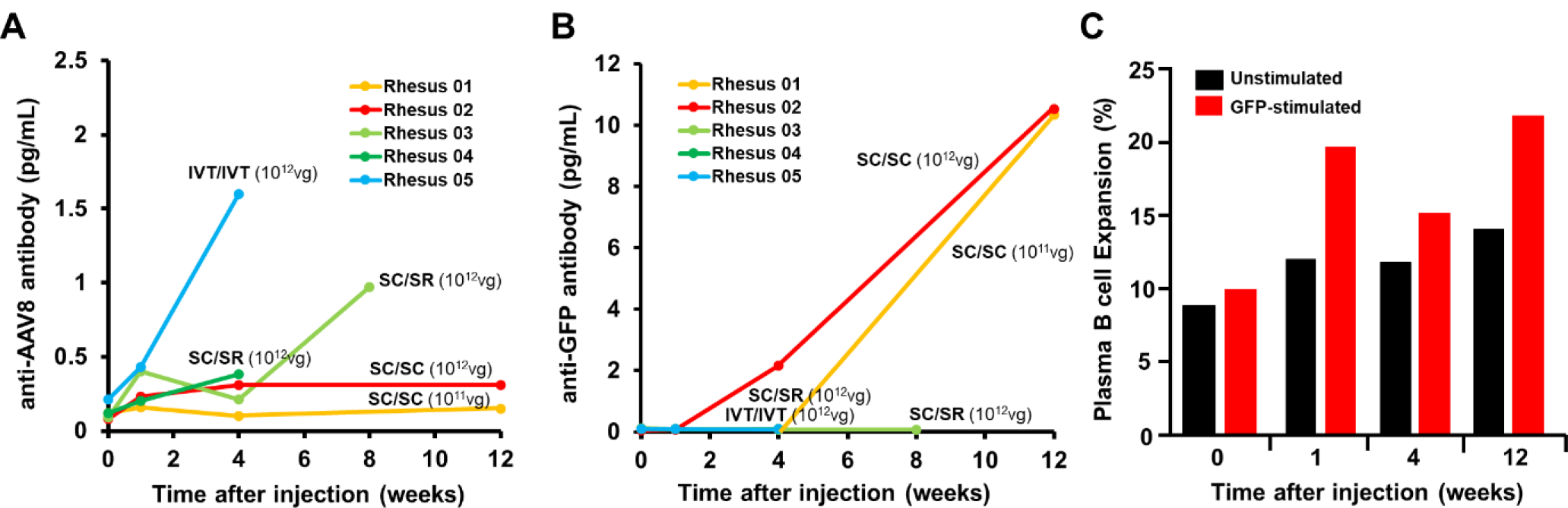
B cell-mediated humoral immune responses against AAV8 and GFP after suprachoroidal injections in NHP eyes. **(A-B)** Line plots compare serum anti-AAV8 antibody (A) and anti-GFP antibody (B) levels in rhesus macaques before and after bilateral suprachoroidal (SC/SC), suprachoroidal / subretinal (SC/SR), or bilateral intravitreal (IVT/IVT) AAV8 injections. **(C)** Bar graphs show flow cytometry analysis of plasma B-cells with expansion upon GFP peptide stimulation from PBMCs collected at various time points after high-dose suprachoroidal AAV8 injections into both eyes in Rhesus 02.

### Cell-mediated immune responses after suprachoroidal AAV8

We next explored cell-mediated immune responses to suprachoroidal AAV8 using ELISpot assays to detect interferon-γ (IFN-γ)-producing T-cells against AAV8 or GFP in PBMCs collected throughout the study and in splenocytes collected at time of necropsy (Figure 4). None of the animals showed appreciable T-cell responses to the AAV8 capsid with the exception of Rhesus 01 which appeared to have pre-existing T-cell responses to AAV8 prior to injection (Figure 4A), despite not having anti-AAV8 antibodies (Figure 3A) or NAbs at baseline.^21^ Similar to the humoral immune responses, suprachoroidal AAV8 also triggered T-cell responses to GFP beginning as early as 1 month after injection, particularly in animals that received suprachoroidal injections in both eyes (Figure 4B). Using splenocytes collected at necropsies, we found suprachoroidal AAV8 injection triggered greater T-cell responses to the GFP transgene product than to the viral vector (Figures 4C-4D, Supplementary Figure 2).

**[Figure 4].**
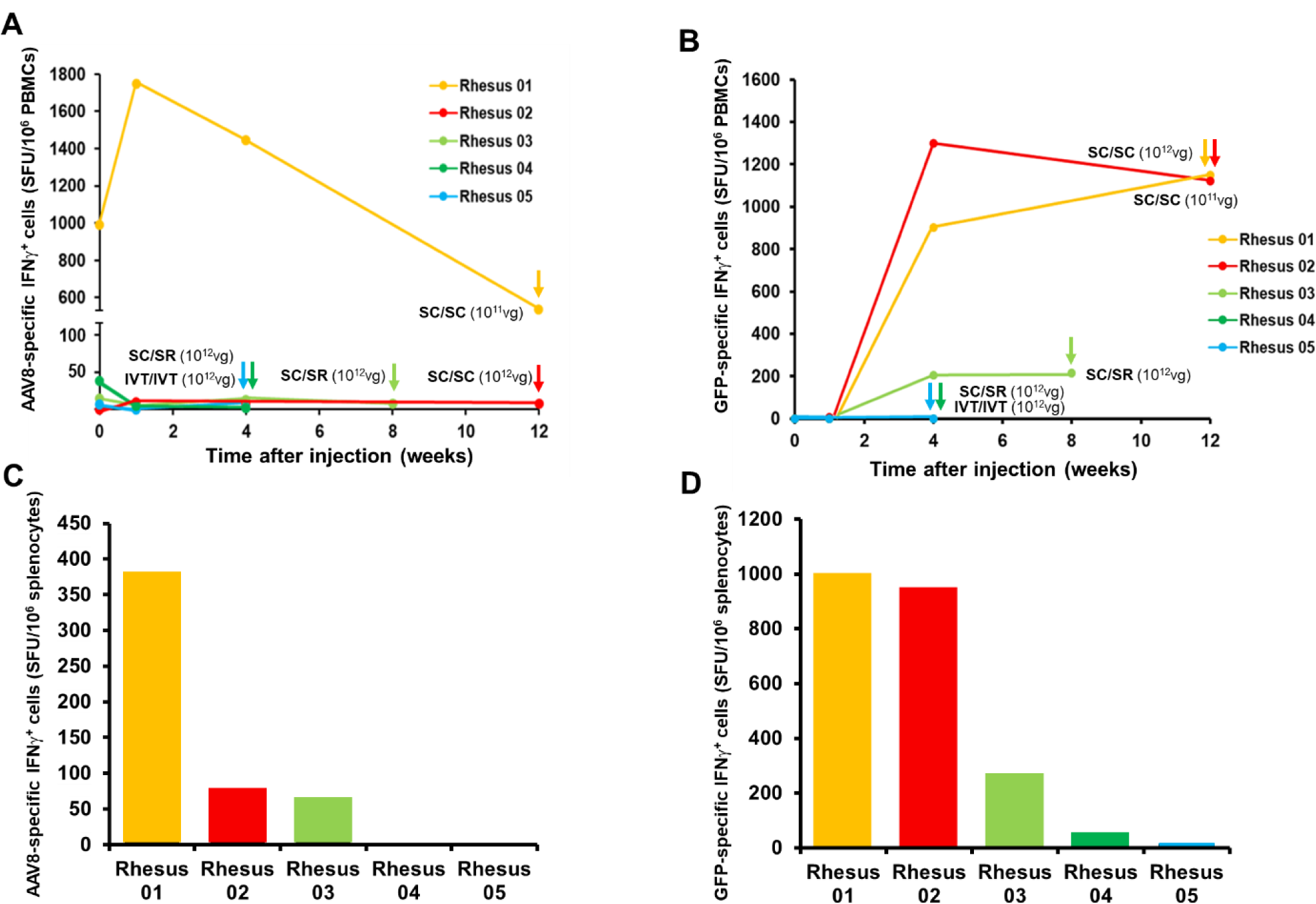
T cell-mediated immune responses against AAV8 and GFP after suprachoroidal injection. **(A-B)** Line plots compare IFN-γ-producing T-cell response against AAV8 capsid (A) and GFP transgene (B) in peripheral blood mononuclear cells (PBMCs) before and after bilateral suprachoroidal (SC/SC), suprachoroidal / subretinal (SC/SR), or bilateral intravitreal (IVT/IVT) AAV8 injections. **(C-D)** Bar plots compare IFN-γ-producing T-cell response against AAV8 capsid (C) and GFP transgene (D) from splenocytes collected at necropsy, as indicated by the corresponding colored arrows for each animal. Abbreviations: SFU, spot-forming units; IFN-γ, interferon gamma

### Systemic biodistribution of suprachoroidal AAV8

To evaluate systemic biodistribution after suprachoroidal AAV8 delivery, we performed quantitative PCR to detect the GFP transgene sequence in genomic DNA from peripheral organs including kidney, liver, and spleen. The highest genome copies were detected in the spleen, followed by the liver, and was undetectable in the kidney (Figure 5). Interestingly, the animal that received intravitreal injection of AAV8 in both eyes (Rhesus 05) showed much higher genome copies of the vector in the spleen and liver, as compared to animals that received suprachoroidal AAV8 in both eyes (Rhesus 01 and 02) or suprachoroidal and subretinal AAV8 in fellow eyes (Rhesus 03 and 04). These studies suggest that suprachoroidal AAV delivery may result in some systemic distribution to peripheral organs such as the spleen and liver, but at much lower amounts than intravitreal injections.

**[Figure 5].**
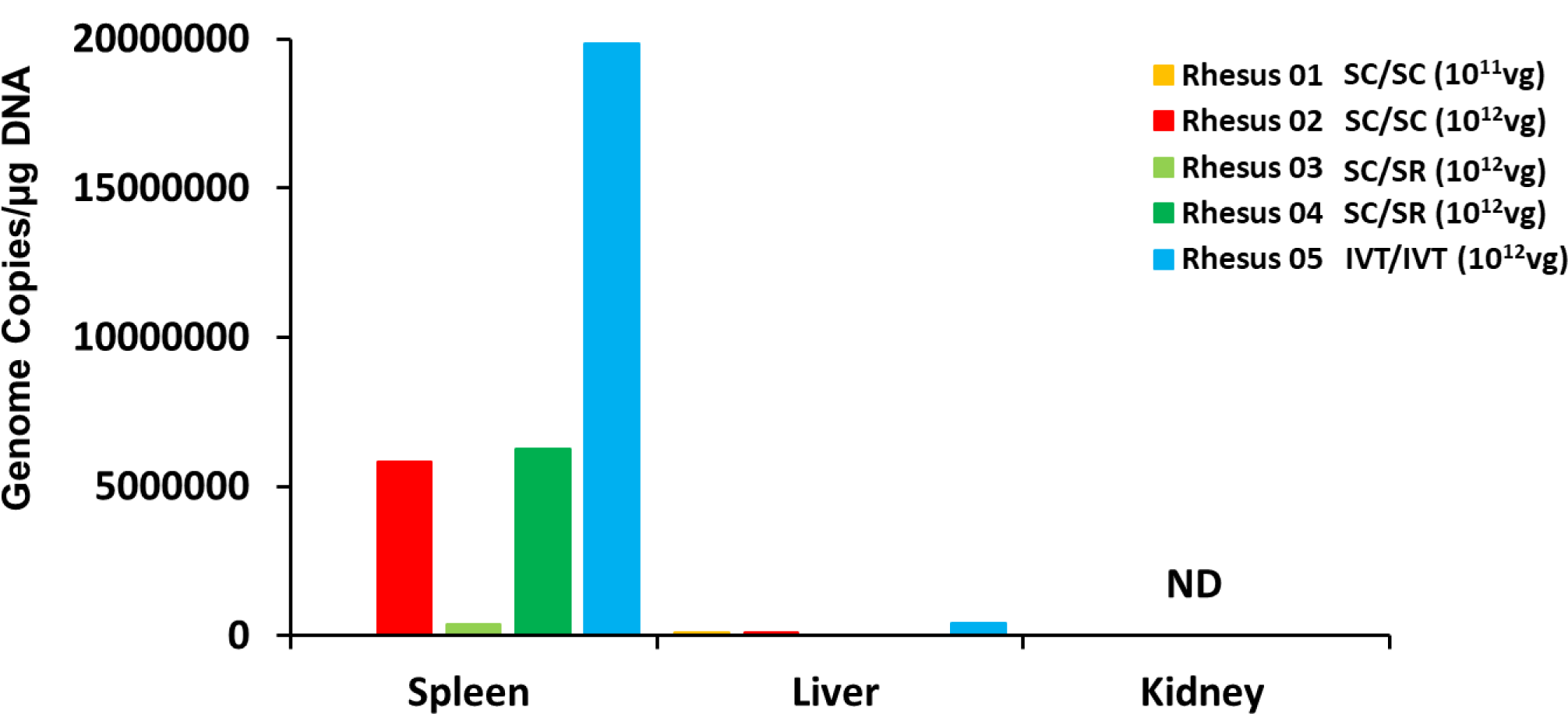
Systemic biodistribution of AAV8 after suprachoroidal injections. Bar graphs show quantification of virally-encoded GFP genome copies measured from peripheral organs including spleen, liver and kidney that were collected at the time of necropsy. Abbreviations: ND, not detected; IVT, intravitreal; SC, suprachoroidal; SR, subretinal.

## Discussion

Despite the presence of ocular immune privilege, AAV-mediated gene delivery to the eye triggers host immune responses that may vary with AAV dose, serotype, route of delivery, and type of transgene. Early studies from the RPE65 gene therapy trials using an AAV2 vector reported a dose-dependent immune response with intraocular inflammation observed in the high-dose (1 × 10^12^ vg/eye), but not low-dose (1 × 10^11^ vg/eye) patient cohorts.^1^ The presence of pre-existing immunity also varies with AAV serotypes, as seroprevalence of anti-AAV2 NAbs in humans has been reported to range between 30-60%, while NAbs against AAV7, AAV8, and AAV9 are lower at 15-30%.^34,35^ Importantly, the route of vector delivery is a major determinant of host immune responses. Intravitreal injections of AAV2 and AAV8 triggers more intraocular inflammation, with more robust humoral and cellular immune responses in mice and NHPs than subretinal delivery,^13,16,36–38^ presumably due to the greater egress of viral particles into systemic circulation from the vitreous cavity. In this study, we evaluated host immune responses to a novel mode of delivering viral vectors into the suprachoroidal space of NHPs using transscleral microneedles.^21,22^ Using an AAV8 vector to express GFP under a CMV promoter, we found that suprachoroidal delivery can trigger a peripheral chorioretinitis and vitritis with outer retinal disruption at month 1 after viral injection, but subsequently showed resolution of inflammation and restoration of retinal architecture at month 3, after systemic corticosteroid administration. The inflammation was accompanied by both humoral and cell-mediated responses to the GFP transgene product, but a less pronounced humoral response to the AAV8 capsid than intravitreal injections.

The host immune responses to the GFP transgene and viral vector can be explained by the pattern of transgene expression, systemic biodistribution of the viral vectors, and unique location of the suprachoroidal space outside the blood-retinal barrier. The blood-retinal barrier is composed of an inner barrier that consists of retinal capillary endothelium, and an outer barrier formed by RPE tight junctions. While the vitreous cavity and subretinal space are immune-privileged ocular compartments within this barrier, the suprachoroidal space is adjacent to the highly-fenestrated choroidal vasculature and readily interfaces with macrophages in the choroid and sclera outside of this barrier. In contrast to intravitreal and subretinal injections which enabled focal GFP transduction within the neurosensory retina, suprachoroidal AAV8 produced broad regions of transgene expression in the RPE and sclera which are outside the blood-retinal barrier. RPE are potent antigen presenting cells (APCs) of the retina,^39,40^ while macrophages and dendritic cells are prevalent in the sclera.^41^ In our study, we observed Iba1+ macrophages/microglia surrounding GFP-expressing RPE, but did not clearly detect any GFP-expressing Iba1+ cells. While the exact cell type responsible for antigen presentation is unclear, our results suggest that immune sensitization likely occurs locally in the eye rather than in peripheral tissues, as both humoral and cellular immune responses to GFP appeared to correlate with the greater transgene expression in the sclera after suprachoroidal injections, regardless of AAV dose, rather than to the higher amounts of viral genomes in peripheral organs after intravitreal delivery. This hypothesis and our results are consistent with the study by Vanderberghe et al, in which T-cell responses against GFP but not AAV capsid were found in NHP eyes after subretinal AAV8 delivery.

Even though the suprachoroidal space is outside the blood-retinal barrier, intravitreal AAV8 triggered a more robust humoral response to the viral capsid, likely due to greater systemic exposure to the AAV8 vector as shown in our biodistribution studies. Trabecular outflow through the canal of Schlemm accounts for 80-90% of vitreous and aqueous humor drainage from the eye, while uveoscleral outflow which likely mediates AAV egress from the suprachoroidal space is less efficient.^42^ Our findings are consistent with previous studies demonstrating greater humoral immune responses after intravitreal versus subretinal injections,^36,37,43^ and suggest that the suprachoroidal space may have better retention of viral particles than the vitreous cavity.

Although the current study focused on AAV8-binding antibodies, we previously found a similar pattern of NAb response that was also more pronounced after intravitreal than suprachoroidal AAV delivery. NAbs prevent viral particles from phagocytosis by blocking essential receptor interactions between the virus and host cells, and may also sequester AAV distribution to the spleen.^44^ By contrast, the role of non-neutralizing antibodies is unclear, and may enhance the clearance of AAV vectors through opsonization or have the opposite effects of NAbs.^44,45^ Interestingly, although serum NAbs can impact the re-administration of AAV given intravitreally,^43^ they do not appear to affect the functional effectiveness of AAV readministered subretinally.^46^ Because a major advantage of suprachoroidal AAV delivery is the capacity for repeated injections, future studies are necessary to determine if the effectiveness of suprachoroidal AAV re-administration may be impacted on repeated dosing.

Our biodistribution assays demonstrated greater peripheral distribution of viral genomes to the spleen and liver after intravitreal injections, compared with suprachoroidal AAV8 delivery, similar to findings by Seitz and colleagues who also found more viral genomes in peripheral organs after intravitreal versus subretinal AAV8 in NHPs.^47^ The higher expression in the spleen alludes to a deviant immune response similar to anterior chamber associated immune deviation (ACAID) – a phenomenon in which immunogen bearing APCs from the eye migrate through the trabecular meshwork to the spleen, where afferent CD4+ Th1 cells and efferent CD8+ cytotoxic T cells differentiate and mature.^15^ Further studies to distinguish more pro-inflammatory from immunosuppressive T-cell subtypes could elucidate the nature of the host cellular immune responses, and help refine strategies for mitigation. The timing of T-cell-directed immunosuppression, for example, has been shown to impact transgene immunogenicity after subretinal AAV delivery.

There are several limitations to our study. Like humans, rhesus macaques are native hosts of AAVs without significant disease association,^48,49^ but exhibit higher seroprevalence of pre-existing immunity to AAV8 capsids.^50,51^ Although we pre-screened animals for the absence of NAbs against AAV8, one animal in our study was found to have a pre-existing T-cell response. Also, the AAV vectors in our study were not generated under Good Medical Practice (GMP) conditions, and may exhibit greater immunogenicity. In addition, although NHPs are excellent preclinical models due to their similar ocular anatomy and immune responses, they do not mount the same level of AAV-specific T-cell responses as humans in liver-directed gene transfer, possibly due to differences in AAV life cycles between humans and NHPs, more efficient recruitment of primed human T-cells to the liver,^52–54^ or loss of inhibitory sialic acid-recognizing Ig superfamily lectins on human T-cells. Finally, because two animals in our study also had subretinal AAV injections in their contralateral eyes, their immune responses may not fully reflect the consequences of suprachoroidal delivery. However, as previous studies have shown that subretinal injections elicit minimal humoral or cellular responses,^13,37^ we believe that the immune responses in these animals are more likely attributable to the suprachoroidal injections.

Suprachoroidal injection of AAV8 is currently under evaluation in human clinical trials for expressing a monoclonal antibody fragment to neutralize vascular endothelial growth factor for treatment of neovascular age-related macular degeneration. Unlike the GFP transgene in our study which is a known immunogen and not native to primate species,^55^ these ongoing human trials employ human-based transgenes and are less likely to generate as robust an immune response. Our study also employed a CMV promoter which has been associated with ocular toxicity not otherwise observed using photoreceptor-specific promoters for AAV transgene expression.^56^ Future studies that employ human-derived and more clinically-relevant promoters and transgenes could better predict host immune responses after suprachoroidal AAV injections, and help facilitate clinical translation of this unique route of vector delivery for retinal gene therapy.

## Materials and Methods

### Animals

The California National Primate Research Center (CNPRC) is accredited by the Association for Assessment and Accreditation of Laboratory Animal Care (AAALAC) International. All studies using rhesus macaques (*Macaca mulatta*) followed the guidelines of the Association for Research in Vision and Ophthalmology (ARVO) Statement for the Use of Animals in Ophthalmic and Vision Research, and complied with the National Institutes of Health (NIH) Guide for the Care and Use of Laboratory Animals. All procedures were conducted under protocols approved by the University of California, Davis Institutional Animal Care and Use Committee (IACUC).

### AAV8 production and intraocular injection

The AAV cis construct which expresses enhanced GFP under a CMV promoter was packaged into AAV8 capsid and purified by the UC Davis NEI Vision Molecular Construct and Packaging Core. After animal sedation, eyes were sterilely prepped with 1% povidone-iodine and flushed with sterile saline, followed by placement of an eyelid speculum. For transcleral microneedle injections, a 700 μm-long 30-gauge microneedle (Clearside Biomedical) was inserted through the conjunctiva and sclera at 4 mm or 10 mm posterior to the corneal limbus to inject into in the superotemporal quadrant (single 100 μL injection) of left eyes, and both superotemporal and inferonasal quadrants (two 50 μL injections) of right eyes. For intravitreal injections, a 0.5 inch-long 30-gauge needle (BD biosciences) was inserted through the pars plana, 4 mm posterior to the limbus, in the inferotemporal quadrant (single 100 μL injection) of both eyes. The viral concentrations are reported in Supplementary Figure 1. Intraocular pressure (IOP) was measured following intraocular injections, and an anterior chamber tap was performed using a 30-gauge needle to remove aqueous until the IOP was normalized.

Rhesus 02 which received high-dose suprachoroidal AAV8 showed signs of ocular irritation and was found to have mild AC cells at 2 weeks after the injection, and was treated with oral prednisone (1mg/kg) for 2 weeks. In Rhesus 03, 04, and 05, a 40 mg periorbital subtenon injection of triamcinolone acetonide suspension (Kenalog-40, Bristol-Myers-Squibb) was also given in the superotemporal quadrant at the request of the veterinarian to prevent uveitis.

### Imaging

All animals underwent SLO and SD-OCT imaging using the Spectralis HRA+OCT device (Heidelberg Engineering, Heidelberg, Germany) before and at 1 week, 1 month, and 2 or 3 months after AAV injections. Confocal SLO was used to capture 55° x 55° or 30° x 30° fluorescence images using 488 nm excitation light and a long-pass barrier filter starting at 500 nm. Images were captured from the central macula and from the peripheral retina by manually steering the Spectralis device. Due to the facial contour of these animals, the superior quadrants could be seen on live visualization but was difficult to capture at sufficient quality for image montage. SD-OCT was performed alongside infrared reflectance images using an 820 nm diode laser to capture 30 ° x 5° SD-OCT raster scans with 1536 A-scans per B-scan and 234 μm spacing between B-scans, in high-resolution mode. SD-OCT scans were captured from the central macula and in regions of visible GFP fluorescence, especially near the junction between transduced and untransduced tissues. 25 scans were averaged for each B-scan, using the Heidelberg eye tracking Automatic Real-Time (ART) software. Animals also underwent color fundus photography (CF-1, Canon) for documentation of clinical exam findings when possible.

### PBMC and splenocyte collection

For PBMC isolation, anticoagulated blood was diluted in phosphate buffered saline (PBS), layered over Ficoll Paque Premium (GE Healthcare, 17544202), and centrifuged for 30 minutes at 800 x g. The PBMC fraction was transferred to PBS and centrifuged again, followed by lysis of red blood cells using Ammonium-Chloride-Potassium (ACK) lysis buffer (Gibco, A1049201), washing with Roswell Park Memorial Institute (RPMI) buffer, and resuspension in 10% dimethyl sulfoxide (DMSO) in heat-inactivated fetal bovine saline (FBS). For splenocyte collection, spleen tissues were homogenized in sterile PBS, passed through a cell strainer, centrifuged, then resuspended in ACK lysis buffer, washed with PBS, and resuspended in 10% DMSO in FBS.

### Binding antibody assay

Binding antibody assays were performed to detect antibodies against GFP and AAV capsid in NHP sera as described previously.^38^ For anti-AAV8 antibody detection, a sandwich-ELISA kit designed for AAV8 titration was used (Progen, PRAAV8). Briefly, microtiter strips with AAV8-specific antibodies were incubated with AAV8 particles (2×10^12^ vg/mL) overnight at 4°C, blocked with 5% milk in PBS, then incubated with macaque sera (1:1000 dilution) at 37°C for 2 hours. After washing, the strips were incubated with horse radish peroxidase (HRP)-conjugated anti-rhesus secondary antibodies (Southern biotech, 6200-50, 1:2000) for 2 additional hours at room temperature, incubated with 3,3’,5,5’-tetramethylbenzidine (TMB; Southern Biotech, 0410-01), stopped with a stopping solution (Southern Biotech, 0412-01), then read with a plate reader (Fisher Scientific accuSkan FC, N16612) with 450 nm absorbance. For detecting anti-GFP antibodies, a 96-well plate was coated with enhanced GFP protein (BioVision, 4999-100, 5 µg/mL) overnight at 4°C, blocked with 5% non-fat milk in PBS, then incubated with diluted serum samples (1:5000) at 37°C for 2 hours followed by detection with HRP-conjugated anti-rhesus IgG as described above. Commercial anti-AAV8 (Progen, 610160S, 1:100) and anti-GFP (Abcam, ab6556, 1:1000) antibodies were used as positive controls, and all values were determined from triplicates. The antibodies were calculated against a standard curve and normalized with total protein.

### Enzyme-linked immune absorbent spot (ELISpot)

ELISpot assays to detect IFN-γ-secreting cells from PBMCs were performed with a commercial kit according to the manufacturer’s instruction (U-CyTech, CT121). Briefly, a 96-well PVDF membrane-bottomed plate was activated with 70% ethanol, and coated with anti-IFN-γ antibodies overnight at 4°C. After washing and blocking, PBMCs were seeded at 4×10^5^ cells per well in RPMI-160 media containing a mix of 182 AAV8 capsid or enhanced GFP peptides (15mers and 11 overlaps, 4 ng/uL, JPT, PM-AAV8-CP, PM-EGFP) for 48 hours. We incubated the cells with Phorbol 12-myristate 13-acetate (PMA, 80 nM) and ionomycin (1.3 µM) for positive control, and DMSO (0.05%) for negative control. After removing the cells, the plate was incubated with biotinylated detection antibody for 2 hours followed by Stretavidin-HRP and 3-Amino-9-ethylcarbazole (AEC) substrate. Spots were counted and normalized with negative control. Spot forming unit (SFU) was calculated from triplicates converted to SFU per 10^6^ cells.

### Quantitative polymerase chain reaction

Systemic biodistribution assays were performed using qPCR with SYBR Green. Liver, spleen and kidney samples were collected at necropsy, and genomic DNA (gDNA) extracted using a commercial kit following the manufacturer’s instruction (Qiagen, 69504). For qPCR, each reaction contained 10 ng of gDNA with SYBR green qPCR master mix (Invitrogen) and forward and reverse primers. qPCR cycling was 95°for 10 min, and 40 cycles of 95°for 10 min, 60°for 1 min, and melting curve analysis was performed for primer dimers. Copy number of GFP transgene was calculated against standard curve, and rhesus beta actin primer set was used as an internal control in a separate reaction. The primer sets used in this study are enhanced GFP forward 5’-AGATCCGCCACAACATCGAGG-3’, GFP reverse 5’-AGCAGGACCATGTGATCGC-3’, beta-actin forward 5’-GGGCCGGACTCGTCATAC-3’ and beta-actin reverse 5’-CCTGGCACCCAGCACAAT-3’. The limit of detection was 162 copies/µg DNA.

### Immunohistochemistry

Immunohistochemistry was performed as described previously.^21^ Posterior eye cups were fixed with 4% paraformaldehyde (PFA) for 2 hours after removal of anterior segments lens and vitreous. After washing with PBS, tissues were cryoprotected with 30% sucrose overnight, then embedded and cryosectioned at 18µm. For antibody labelling, sections were washed with PBS, blocked with 10% normal donkey serum for 30 min, then incubated in primary antibody for 1-2 hours at room temperature, followed by Alexa Fluor-conjugated secondary antibodies. Primary antibodies include IBA-1 (Wako, AB10558, 1:100), GFAP (Dako, Z0334, 1:200), and CD45 (BD, 552566, 2.5µg/ml).

### Flowcytometry

For flow cytometry, 0.5 × 10^6^ PBMCs or splenocytes per well were plated in duplicate in 96-well plates in RPMI supplemented with 10% FBS for 24 hours. The cells were stimulated with AAV8 peptide (4ng/μl, JPT, PM-AAV8-CP), GFP peptide (4ng/μl, JPT, PM-EGFP), cRPMI alone (unstimulated), or with PMA (80 nM)-Ionomycin(1.3 μM) (positive control). Cultures were incubated at 37°C for 48 hours, washed with PBS, and stained for flow cytometric analysis. The cells were incubated with 50µL of an antibody cocktail for CD8 (Thermo Fisher, Q10055), HLADR (BioLegend, 307656), CD19 (BioLegend, 302239), CD27 (Biolegend, 302824), and CD38 (Labcome, 100825) for 30 minutes at room temperature in the dark, followed by 2 washes with FACS buffer (PBS + 1% FBS), and resuspended in 300 µL of FACS buffer for analysis. The data were acquired within an hour on a BD FACS LSR II flow cytometer (Beckman Coulter Life Sciences, USA).

## Acknowledgements

We thank Marie Burns and Huaiyang Chen for AAV production, Amanda Carpenter for PBMC and splenocyte preparation, Jeffrey Roberts and John Morrison for CNPRC logistics, Monica Motta for assistance with animal imaging, and Dennis Hartigan-O’Connor for discussions. We also thank Clearside Biomedical and Glenn Noronha, Jesse Yoo, and Donna Taraborelli for providing the suprachoroidal microneedles used in our study. This study was supported by the California National Primate Research Center pilot grant program and base grant NIH P510D011107. GY is supported by NIH K08 EY026101, NIH R21 EY031108, and Macula Society. ST is supported by U24 U24EY029904. The AAV8 vector was produced by the Center for Vision Sciences Molecular Constructs and Packaging core facility supported by NIH P30 EY012576. No funding organizations had any role in the design or conduct of this research. The content is solely the responsibility of the authors and does not necessarily represent the official views of the funding agencies.

## Author Contributions

G.Y conceived the study design, obtained funding, performed the examinations, intraocular injections, and in vivo imaging, and supervised the study. S.H.C, T.S, and T.N conducted the binding antibody assays, ELIspot assays, biodistribution studies, and immunohistochemistry. I.M assisted clinical examination, injections, and tissue collection. A.M conducted flow cytometry and analyzed data. G.Y and S.H.C analyzed all data and wrote the manuscript. G.Y, S.H.C, T.C, P.S and S.T critically reviewed the manuscript.

## Figure Legends

**[Supplementary Table 1].** Summary of study animals, demographic, injection mode, dose and necropsy dates. Abbreviations: OD, right eye; OS, left eye; IVT, intravitreal; SC, suprachoroidal; SR, subretinal; vg, viral genomes

| Animal ID | Age<br>(Years) | Sex | Injection Mode<br>(OD/OS) | Viral Dose<br>(vg/eye) | Necropsy<br>Date |
| --- | --- | --- | --- | --- | --- |
| Rhesus 01 | 10.25 | Female | SC/SC | 7x10<sup>11</sup> vg | Month 3 |
| Rhesus 02 | 9.05 | Female | SC/SC | 7x10<sup>12</sup> vg | Month 3 |
| Rhesus 03 | 5.47 | Male | SR/SC | 7x10<sup>12</sup> vg | Month 2 |
| Rhesus 04 | 9.38 | Female | SR/SC | 7x10<sup>12</sup> vg | Month 1 |
| Rhesus 05 | 10.38 | Male | IVT/IVT | 7x10<sup>12</sup> vg | Month 1 |

**[Supplementary Figure 1].**
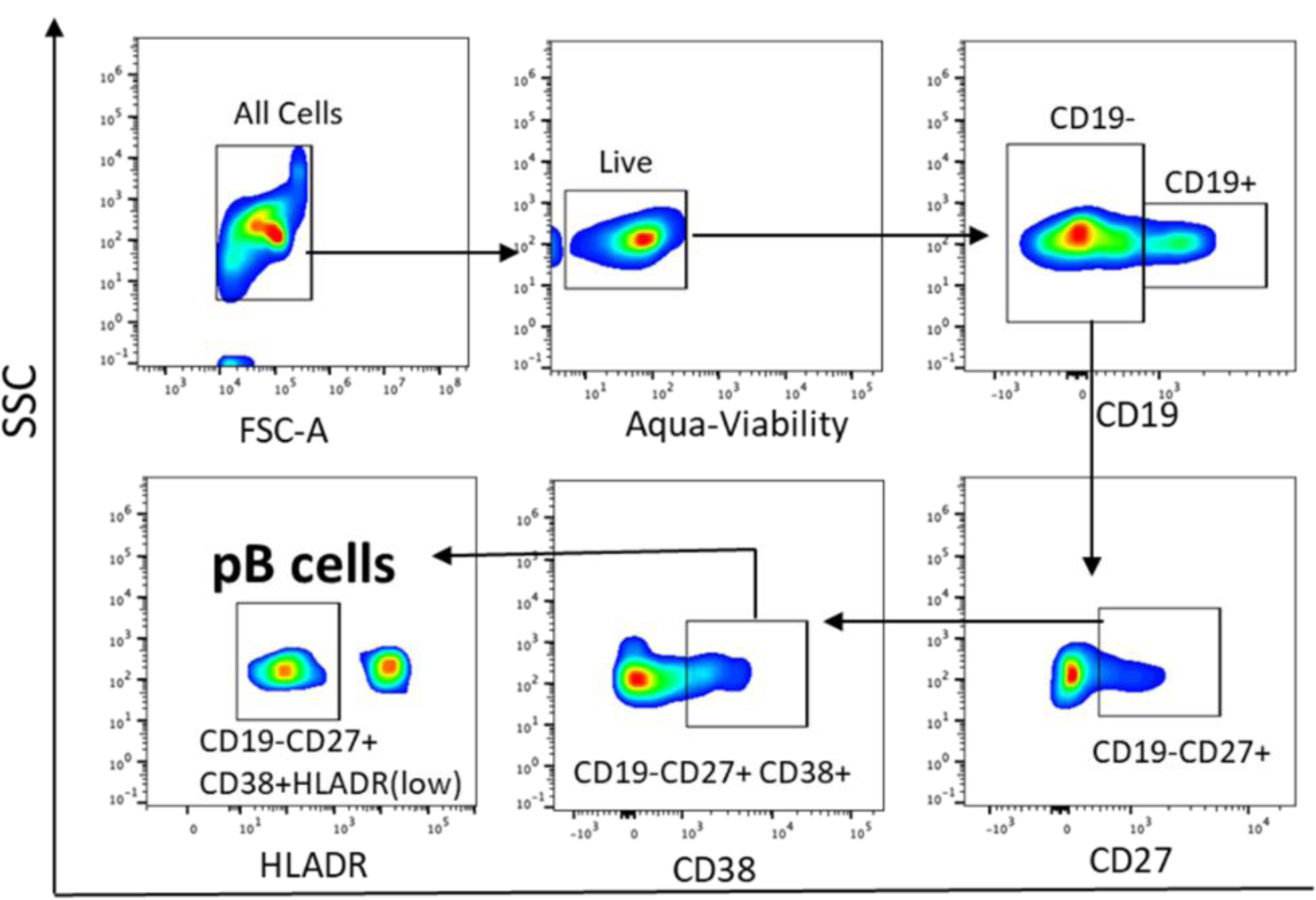
Flow cytometry gating strategy for plasma B-cell quantification.

