## Supplementary figures and images for "Host immune responses after suprachoroidal delivery of AAV8 in nonhuman primate eyes"

### Supplementary Figure 1

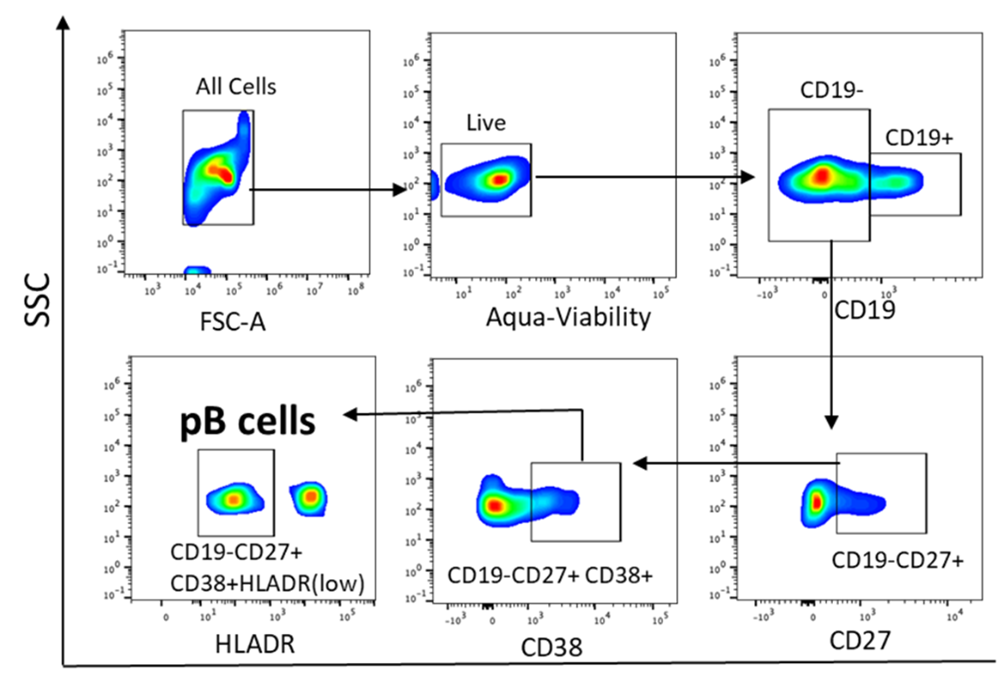

### supplementary Table 1

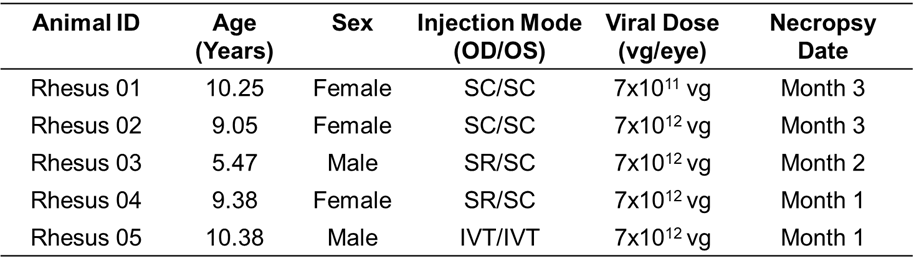
